# Data availability of open T-cell receptor repertoire data, a systematic assessment

**DOI:** 10.1101/2022.04.13.488243

**Authors:** Yu-Ning Huang, Naresh Amrat Patel, Jay Himanshu Mehta, Srishti Ginjala, Petter Brodin, Clive M Gray, Yesha M Patel, Lindsay G. Cowell, Amanda M. Burkhardt, Serghei Mangul

**Affiliations:** Department of Clinical Pharmacy, School of Pharmacy, University of Southern California, Los Angeles, California, 90089, USA; Department of Pharmaceutical Sciences, School of Pharmacy, University of Southern California, Los Angeles, California, 90089, USA; School of Computing and Electrical Engineering, Indian Institute of Technology, Mandi, Himachal Pradesh, India; Department of Immunology and Inflammation, Imperial College London, United Kingdom; Department of Women’s and Children’s Health, Karolinska Institutet, Sweden; Division of Molecular Biology and Human Genetics, Biomedical Research Institute, Stellenbosch University, Cape Town, South Africa; Department of Clinical Pharmacy, School of Pharmacy, University of Southern California, 1540 Alcazar Street, Los Angeles, CA 90033, USA; Division of Biomedical Informatics, Department of Population and Data Sciences, Department of Immunology, University of Texas Southwestern Medical Center at Dallas, USA; Department of Clinical Pharmacy, School of Pharmacy, University of Southern California, CA, USA; Department of Clinical Pharmacy, School of Pharmacy, University of Southern California, CA, USA,, Twitter: @smangul1

## Abstract

The improvement of next-generation sequencing technologies has promoted the field of immunogenetics and produced numerous immunogenomics data. Modern data-driven research has the power to promote novel biomedical discoveries through secondary analysis of such data. Therefore, it is important to ensure data-driven research with great reproducibility and robustness for promoting a precise and accurate secondary analysis of the immunogenomics data. In scientific research, rigorous conduct in designing and conducting experiments is needed, specifically in scientific and articulate writing, reporting and interpreting results. It is also crucial to make raw data available, discoverable, and well described or annotated in order to promote future re-analysis of the data. In order to assess the data availability of published T cell receptor (TCR) repertoire data, we examined 11,918 TCR-Seq samples corresponding to 134 TCR-Seq studies ranging from 2006 to 2022. Among the 134 studies, only 38.1% had publicly available raw TCR-Seq data shared in public repositories. We also found a statistically significant association between the presence of data availability statements and the increase in raw data availability (p=0.014). Yet, 46.8% of studies with data availability statements failed to share the raw TCR-Seq data. There is a pressing need for the biomedical community to increase awareness of the importance of promoting raw data availability in scientific research and take immediate action to improve its raw data availability enabling cost-effective secondary analysis of existing immunogenomics data by the larger scientific community.

## Introduction

Advanced high throughput sequencing technologies have reshaped the landscape of contemporary immunology research providing researchers with a rich set of tools and methods to study fundamental aspects of immune responses across a variety of disciplines. Raw TCR-Seq data is produced accompanying the development and progress of immunogenomics research. Availability of raw TCR-Seq data allows effective re-analysis of the data^1^ (also known as secondary analysis) which in turn may accelerate novel biomedical discoveries^2^. Using public repositories to share raw data substantially simplifies discovering and retrieving datasets of interest, as one has scalable, programmatic access to a large number of studies. Over the last decade, there has been tremendous progress to improve sharing of raw immunogenomics data, which allows researchers to easily obtain the various types of data stored in public repositories^3^ (e.g. SRA^4^). But to unlock the full potential of publicly available raw TCR-Seq data for the secondary analyses, it is crucial for a study to be conducted accurately with reproducible and reliable laboratory practices and accurate results^5^. Moreover, the research should report publicly available raw data accompanying metadata for secondary analysis^6,7^. One of the merits of science is to be able to leverage published data for subsequent novel scientific discovery^8^. To ensure the results are reproducible and have sufficient quality for accurate and precise secondary analysis, the data should be shared under the FAIR principle–Findability, Accessibility, Interoperability, and Reusability–to ensure the quality of the raw data to be reused accurately and efficiently in research^8,9^. Scientific communities, peer-reviewed journals, funding agencies, and government agencies have emphasized the importance of making raw omics data publicly available with free access to the public^7^. As such, establishing protocols, guidelines, and authors’ checklists for reporting the raw immunogenomics data in research, and peer-review journals mandating authors to upload the data to public repositories are potential means to increase the raw data availability^3,7,10^.

It is also crucial to ensure studies share the raw data used for publication and analyses. In the fields of immunogenetics studies since the development of next-generation sequencing tools promotes research on genetic sequencing and production of raw immunongenomics data due to the reduction in price for genetic sequencing^11^. T cell receptor sequencing (TCR-Seq) studies allow researchers to profile human immune systems providing updated insights into T cell receptors. TCR-Seq data allows researchers to examine a unique individual’s immune status, immune responses, and compare the T cell populations between individuals in healthy or disease states, such as autoimmune diseases, infectious diseases, and cancer^12–14^. Additionally, TCR-Seq studies allow the development of novel therapeutics and biomarkers, including diagnostics^15^ for autoimmune disease^16^ and cancer^17–19^, CAR-T cell therapy^20^, vaccines^21^, and monoclonal and therapeutic antibodies^22^. Due to the production of vast TCR-Seq data in the field of TCR repertoire studies, there is a pressuring need to ensure that raw sequencing data from TCR-Seq studies is provided with full access to the public to facilitate further computational and bioinformatics research in the field. In this study, we examined TCR-Seq studies for the availability of the raw sequencing data across 134 TCR-Seq studies over 11,918 samples and examined the public genomic repositories where the researchers stored the TCR-Seq data from their analyses. We found, only 38.1% of TCR-Seq studies have made the overall raw data available, and that the majority of studies (35 studies) with available raw data stored the raw data in Sequence Read Archive^4^. Further, 61.9% of the TCR-Seq studies did not share the raw data in the original publications or that the raw data will only be available upon request. We discussed the potential barriers researchers are facing that deter them from sharing the raw data and the potential ways to improve the availability of the raw data in the field of TCR-Seq studies.

### The availability of raw sequencing data in TCR-Seq studies is limited

We investigated 134 published TCR-Seq studies across 11,918 samples in PubMed for the raw sequencing data availability ranging from 2006 to 2022. The studies were considered as having available raw sequencing data if we could acquire the samples of the studies’ raw FASTQ or FASTA files. According to our results, only 38.1% (51 out of 134 studies) of the TCR-Seq studies shared raw TCR-Seq data in the original publications at public genomic repositories (Figure 1a). Conversely, 61.9% (83 out of 134 studies) of the TCR-Seq studies have unavailable raw RNA-Seq data or have raw data available upon request (Figure 1a). We observed a similar trend of raw data availability among the 11,918 samples of the 134 TCR-Seq studies, in which 25% of the samples among the TCR-Seq studies had available raw data (Figure 1a). We also observed that the raw TCR-Seq data availability has increased over the past decade (Figure 1b). Among the 134 studies, 89.6% (120 studies) of the raw data were generated in medical research institutes, including medical schools, hospitals, medical centers, private health/disease research institutes, and government health research institutes, such as the National Institutes of Health (Figure 1c). The remaining raw data were generated in engineering schools and other research institutes (Figure 1c). Additionally, we examined the types of TCR-Seq chains in the available raw sequencing data. Among the samples with available raw sequencing data, 65.2% of the TCR-Seq data were TCR beta (TCRβ) chain, 30.7% of the TCR-Seq data were TCR alpha (TCRα) chain, and 4.1% of the TCR-Seq data were TCR gamma (TCRγ) or TCR delta (TCRδ) chain (Figure 1d).

**Figure 1.**
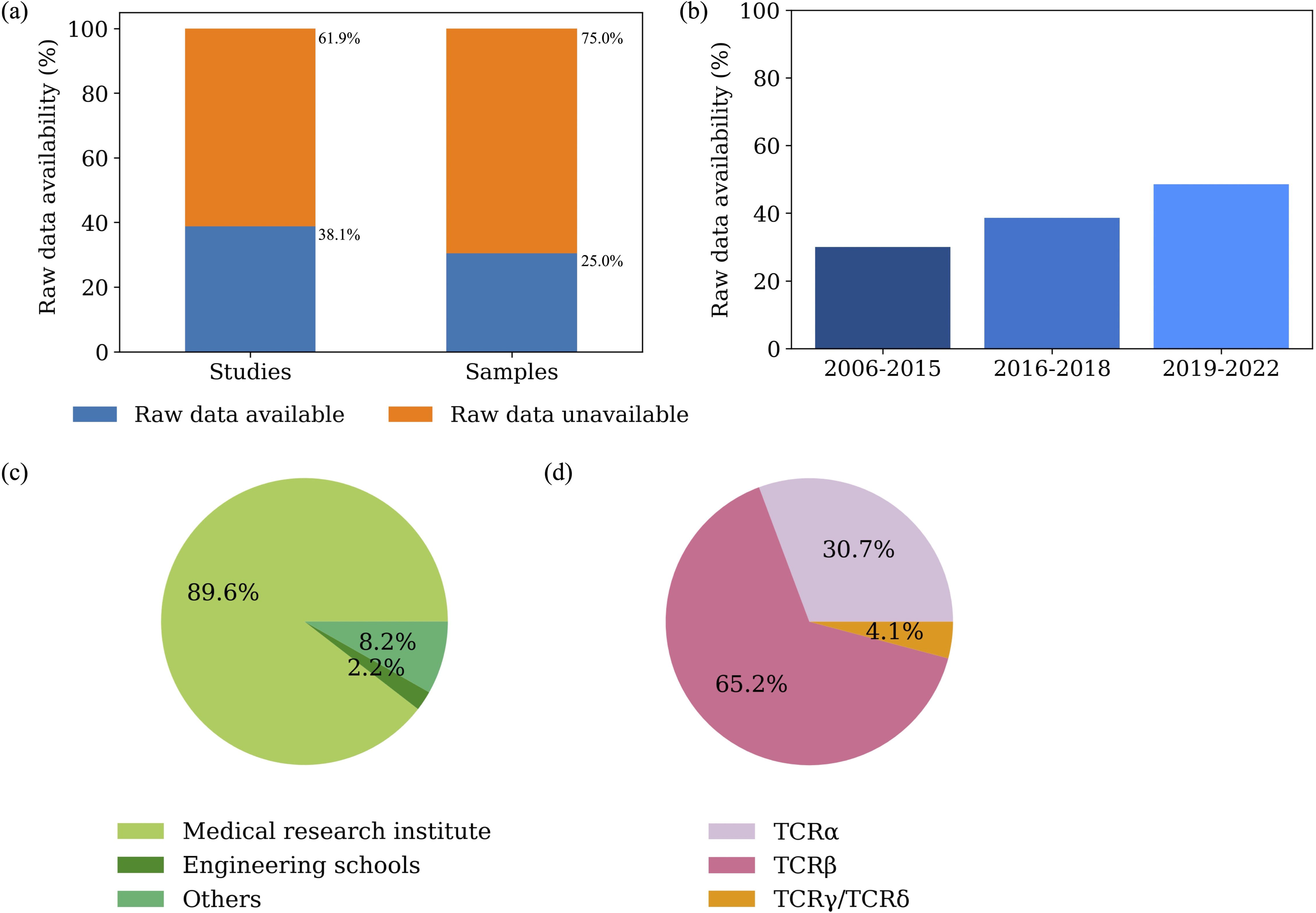
The raw data availability across the 134 TCR-Seq studies. a, The proportion of the raw sequencing data available in the TCR-Seq studies. Left: The raw sequencing data’s availability of the 134 TCR-Seq studies; Right: The raw sequencing data’s availability of the 11,918 samples in the 134 TCR-Seq studies. b, Bar plot of the changes in the raw TCR-Seq data availability from 2006 to 2022. c, The pie chart depicting the types of institutes where the data was generated. d, The pie chart depicting the types of the chain types of TCR-Seq data.

We further investigated the specific reasons that the raw data was unavailable among the 83 studies without available raw TCR-Seq data. Among the 83 studies with unavailable raw data, rather than sharing raw sequencing data, 44 studies only shared summary data, such as summary data on ImmuneACCESS® (Adaptive Biotechnologies), VDJdb, and supplementary files. Other 34 studies did not include statements about the raw data or provide raw sequencing data in the original publications and five studies indicated that the raw sequencing data will be available upon making direct requests to the authors (Figure 2a).

**Figure 2.**
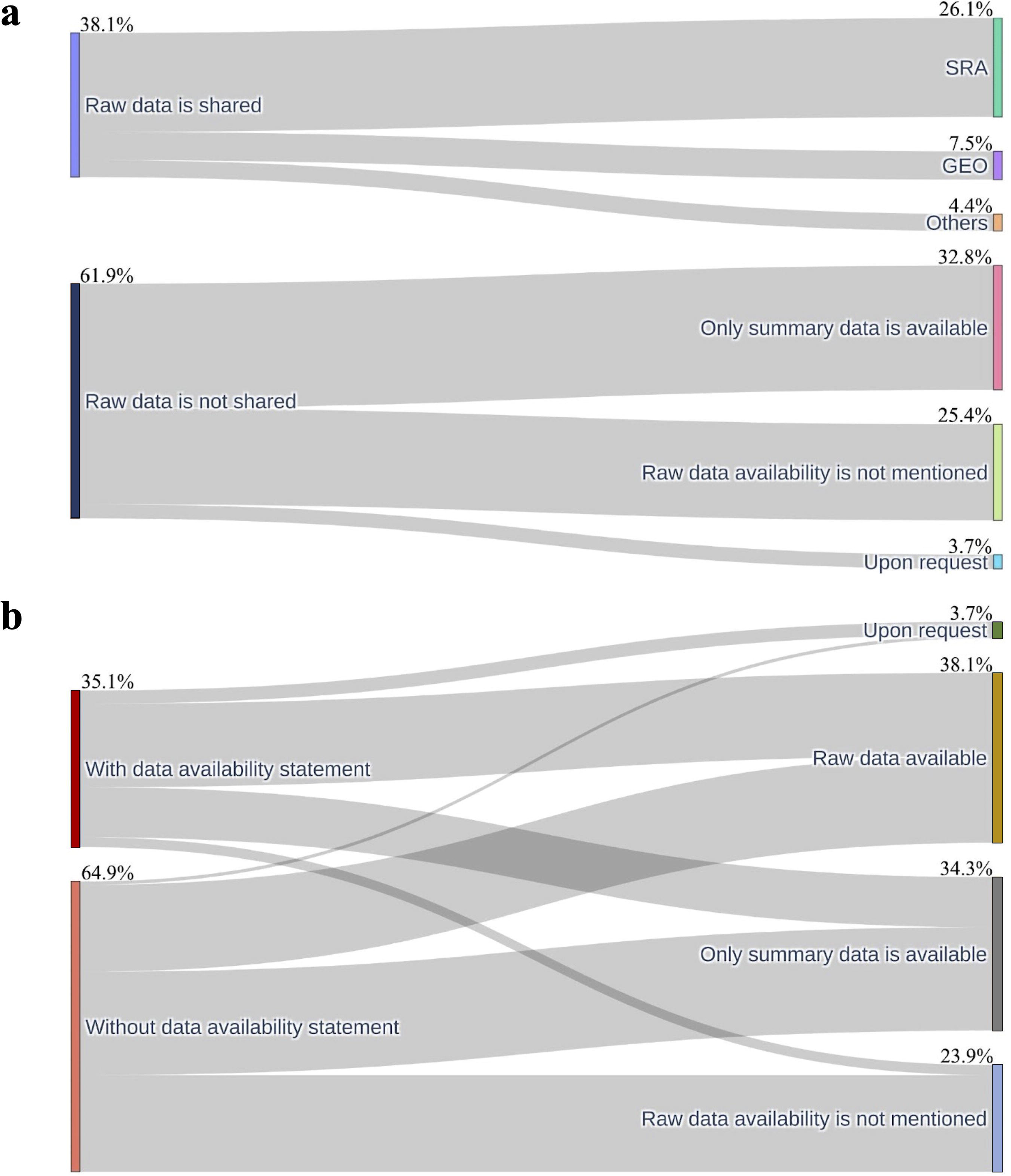
The Sankey plot of the raw data availability across the 134 TCR-Seq studies. a, The Sankey plot depicting the proportion of raw data availability (left) and platforms where available raw data is stored and the specific reasons why the raw data is unavailable (right). b, The Sankey plot shows the proportion of TCR-Seq studies with available “Data availability statements” in the text of the article (left) and the proportion of the studies with available raw data, studies reporting only summary data, and studies mentioning that data is available upon request, and studies not mentioning data availability information.

### The platforms used to store the raw TCR-Seq data

We further examined the platforms researchers used to share raw TCR-Seq data. Among the TCR-Seq studies that shared the raw sequencing data, 35 of the 51 studies (68.6%) of the raw data are shared in Sequence Read Archive (SRA)^4^ and ten of the 51 studies (19.6%) shared the raw data in other repositories, including Gene Expression Omnibus^23^ (GEO). The remaining studies shared the raw sequencing data in various online repositories, including European Genome-phenome Archive^24^ (EGA), and National Genomics Data Center, China^25^ (NGDC) (Figure 2a).

### The presence of the “Data availability statement” increases the availability of raw TCR-Seq data

As part of promoting more transparent and reproducible research, many journals do require data availability statements for studies to be published in the journals^26,27^. We examined the impact of the presence of the “Data availability statement” in research articles on the availability of raw TCR-Seq data. According to the results, the presence of data availability statements improves the raw data availability by 23.3% (26 out of 87 studies without data availability statements) to 53.2% (25 out of 47 studies with data availability statements) (Supplementary table 1).

There are 47 studies containing “Data availability statements” in the corresponding publications. Among the 47 studies with “Data availability statements”, 53.2% (25 out of the 47 studies) studies shared the raw TCR-Seq data in the publications while the rest of the studies did not share raw TCR-Seq data in the original publications or would be available upon request. Conversely, there are 87 studies that did not have “Data availability statements” in the original publications. For the 87 publications that do not have the “Data availability statement”, only 29.9% (26 out of the 87 studies) of the studies shared the raw TCR-Seq data in the publications. There are three studies with data availability statements but provide unavailable accession numbers, so we cannot access the raw data of the three studies. Therefore, we categorized them into the studies with available data availability statements and raw data availability is not mentioned. We conducted a Pearson’s chi-squared test (χ^2^) to examine the associations between the presence of data availability statements and the availability of raw data and found a statistically significant relationship between the two(p=0.014) - the data availability statements are improving access to raw data availability.

We also examined parts of the publications where the raw data availability is mentioned. Among the studies that shared the raw TCR-Seq data, authors included the data availability statements in various locations of the research articles. 49.0% (25 out of 51 studies) had the data availability statements in the “Data availability statement” portion of the research articles. Eight out of 51 studies (15.7%) had the data availability statements in the footnote of the studies, whereas the remaining 18 out of 51 studies (35.3%) mentioned their raw data availability directly in the main text of the studies.

## Discussion

Our study is the first to assess the raw data availability of TCR-Seq studies. According to the results, the raw data availability is alarmingly low, with only 38.1% of the TCR-Seq studies sharing the raw data (Figure 2b). We investigated the association between the presence of data availability statements in the articles and the raw data availability. We discovered that the data availability statements presented have a statistically significant association with the raw data availability (p=0.014). The presence of data availability statements improves the raw data availability by 23.3%. While data availability statements help to improve data availability, we noticed that it is still possible to have such a statement and not share raw data or only share summary data or only will be available upon request (46.8%) (Supplementary table 1). Some studies only shared the summary data in the data availability statement section of the publications. Additionally, three of the studies shared an erroneous SRA accession number in the data availability statement sections of the publications so we were not able to access the raw data of the articles.

According to our analysis, the primary reason for unavailable raw data is that the studies did not indicate the statements of raw data in the research articles nor shared the raw data in the article (34 studies), which did not share the raw sequencing data of the analyses. Secondly, most authors only shared summary data in the original publications (44 studies) reckoning that they shared the data in their research. For example, 23 studies shared the summary data in the corporate-owned repository, ImmuneACCESS® by Adaptive Biotechnologies. Unfortunately, the ImmuneACCESS® repository only offers access to the summary data of the studies, thus the raw data files are still inaccessible to the public, making the researchers unable to re-analyze the raw data generated by Adaptive Biotechnologies for novel biomedical discoveries. Thirdly, several studies mentioned that the data will only be available by making direct requests of the data from the authors (5 studies). However, it has been previously shown that the statements mentioning that the raw data will be available upon request did not guarantee data availability and instead were not a sustainable and practical way to improve research reproducibility and raw data availability^26,27^. The specific reasons for not sharing raw TCR-Seq data are beyond the scope of this manuscript and need to be investigated in future studies. The perceptual and technical barriers researchers are facing when sharing the data are yet to be determined. Previous work has suggested that the reasons for authors to not share the raw data might be because they are unaware of the importance of sharing the raw data, there may be culturally related or technical barriers that deter or prevent authors from sharing the raw data^27^.

Individual researchers, research institutes, and journals should all take part in ensuring raw data availability^27^. Individual researchers and research institutes more specifically should hold the burden of making the raw data publicly available. Journals also have a critical role in promoting raw data availability, and many journals are already taking actions to promote data availability via data availability statement requests within articles upon publication. However, more stringent measures need to be imposed by the journals to ensure raw data availability^28^. Since existing policies for making the data availability statement mandatory in the article are improving raw data availability, our results suggest that journal policies might have the largest potential impact on improving raw data availability. It is known that many journals have already mandated the authors to share the raw data of the studies^29^. Journal’s policies in mandating authors to share the raw data might be a feasible way to improve the raw data availability^30^. In conclusion, the pressing need to increase awareness of enhancing raw data availability in scientific research can enable cost-effective secondary analysis of existing immunogenomics data for novel biomedical discoveries. Therefore, we recommend that all members of the biomedical community, including individual researchers, research institutions, and journals, should contribute to increasing the awareness of raw data sharing and improve the raw data availability in future studies.

