## Supplementary information for "Data availability of open T-cell receptor repertoire data, a systematic assessment"

**Supplementary Material**

**Method**

We collected TCR-Seq studies from PubMed and examined the raw data availability, reasons for not sharing the raw data, and the last authors' affiliations. We collected 134 TCR-Seq studies from PubMed ranging from 2006 to 2022 across 11,918 samples. We examined the availability of raw TCR-Seq data, which the studies will only be considered with available raw data if they shared the raw FASTA or FASTQ files of the studies. We investigated the reasons for studies that do not share the raw data, and categorized the studies into the following three categories, including only summary data is available (meaning that the studies only shared summary data in the publications or shared the summary data on ImmuneACCESS® (Adaptive Biotechnologies), VDJdb, and in the supplementary files), raw data availability is not mentioned (meaning that the studies did not mention the access to the raw data of the studies in the articles, and additionally, the studies that provided falsifying accession numbers making us unavailable to access the raw data of the studie), and upon request (meaning that the raw data will be available by making direct requests to the authors of the publications). The above three categories are reckoned as studies that have unavailable raw data.

We also examined the last author's affiliation and categorized the last authors' affiliations into one of the three categories, medical research institutes, including affiliations with medical schools, schools of medicine, hospitals, medical centers, private health/disease research institutes, and government health research institutes, engineering schools, and others, including other science-related research institutes.

**Table 1.** The number of studies with data availability statements that shared raw data, only summary data is available (shared only summary data), upon request (making data available upon request), and raw data availability is not mentioned (not mentioned any raw data availability statements in the articles); and the number of studies without data availability statements that shared raw data, only summary data is available (shared only summary data), upon request (making data available upon request), and raw data availability is not mentioned (not mentioned any raw data availability statements in the articles).

| **Category (N=47 (with data availability statements); N=87 (without data availability statements))** | **Subcategory** | **Number of studies** |
|---|---|---|
| Studies with data availability statement | Raw data available | 25 |
| | Only summary data is available | 15 |
| | Upon request | 4 |
| | Raw data availability is not mentioned | 3 |
| Studies without data availability statements | Raw data available | 26 |
| | Only summary data is available | 31 |
| | Upon request | 1 |
| | Raw data availability is not mentioned | 29 |
